# Patterns of host plant use by monarch butterflies revealed through annotation of more than 35,000 community science records

**DOI:** 10.64898/2026.04.06.716508

**Authors:** Micah Freedman

**Affiliations:** Department of Ecology & Evolutionary Biology, University of Toronto

**Keywords:** community science, species interactions, plant-herbivore, monarch butterfly, milkweed, iNaturalist, migration, host breadth

## Abstract

Community science data are increasingly recognized as important resources for biodiversity research, in part because of the spatial and temporal resolution that they afford. While these data are useful for applications such as describing occurrence patterns, tracking movement of migratory animals, and recording phenological events, they can also be probed for “second-order” purposes, such as documenting species interactions. Here, I present a dataset of more than 35,000 annotated interactions between monarch butterfly (*Danaus plexippus*) larvae and their associated host plants from the community science platform iNaturalist. I document more than 70 unique species of milkweed hosts (Apocynaceae: Asclepiadoideae) used by monarch larvae, including a number of previously undocumented interactions. Monarchs show strong seasonal turnover in the species of host plants used across the migratory cycle, highlighting the importance of early season hosts like *Asclepias viridis* and *A. asperula* in eastern North America and *A. californica* and *A. cordifolia* in the west. I also demonstrate that non-native horticultural milkweed species have increased the spatial extent of monarch breeding during winter (November – February) by more than 60%, a pattern previously suggested from observational data but not formally quantified until now. To my knowledge, this represents the largest analysis to date of species interactions using unstructured community science data and highlights the value of platforms like iNaturalist for conducting fundamental research in ecology and conservation.

## Introduction

Over recent years, community science platforms—notably including Zooniverse, eBird, iNaturalist, and several others—have become valuable tools in biodiversity research. Of the records archived each year by the Global Biodiversity Information Facility (GBIF), iNaturalist alone now contributes more than 50% of reptiles, amphibians, arachnids, and gastropods, and more than 30% of plants and insects (Mason et al. 2025). By far the most common application of community science-derived data is using occurrence data to understand species’ distributions (Mason et al. 2025). However, in the case of platforms like iNaturalist, in which users upload images to accompany their observations, additional information can be gleaned by analyzing visual features of focal taxa and their surroundings. For example, iNaturalist data have been used in research on plant phenology (Tourville et al. 2024, Mackenzie et al. 2025), animal behavior (Vardi et al. 2021), intraspecific trait variation (Jansen et al. 2024), and description of new species (Rosa et al. 2022).

An emerging use of community science data is the documentation of species interactions (Callaghan et al. 2021). Since 2019, at least 50 studies have used community science data for this purpose (Table S1). Notable examples include studies on the prevalence of ticks on alligator lizards (Putman et al. 2021); patterns of specificity exhibited by bot flies in selecting primate hosts (Ortiz-Zárate et al. 2024); numerous studies on patterns of floral visitation (e.g., Pernat et al. 2024) and plant-frugivore interactions (e.g., Silva et al. 2024); and associations between mistletoes and their host plants (Santiago-Rosario et al. 2024). Community science observations of species interactions can be especially valuable for understanding the biology of taxa that are geographically widespread (e.g., de Groot et al. 2024), for which concerted range-wide sampling might otherwise be challenging, or taxa that are difficult to observe because of their rarity or ephemeral nature (e.g., Fourreau et al. 2024).

Monarch butterflies (*Danaus plexippus*) are a widespread species with a long history of research through community science-based initiatives, including the Monarch Larva Monitoring Project, Journey North, Mission Monarch, and broader efforts like the North American Butterfly Association’s annual counts (Oberhauser et al. 2015; Ries & Oberhauser 2015). These efforts primarily focus on generating data on year-to-year monarch abundance and documentation of migration-related events, such as date of first sightings or aggregations of migratory adults during autumn. Data from iNaturalist, which features more than 440,000 observations of monarchs as of March 2026, are collected in an unstructured way that makes them challenging to use for studying patterns of abundance (Johnston et al. 2021). However, the sheer scale of observations, as well as the associated metadata and information contained within associated images, makes iNaturalist a valuable data source for drawing inferences about interactions between monarchs and their environment.

The more than 108,000 observations of monarch larvae posted to iNaturalist generally include identifying features of monarch host plants (Figure 1), which can be annotated to describe patterns of host use. In this paper, I present what is, to my knowledge, the largest analysis to date of species interaction data sourced from community science images (Table S1) to understand associations between monarchs and their milkweed hosts. In particular, I focus on three specific objectives: (1) Documenting the host breadth of monarchs, both within the genus *Asclepias* and across the broader Apocynaceae; (2) Describing patterns of seasonal turnover in host use and determining which hosts are most important across the migratory cycle; (3) Quantifying the extent to which cultivation of non-native milkweed hosts has affected monarch breeding phenology.

**Figure 1.**
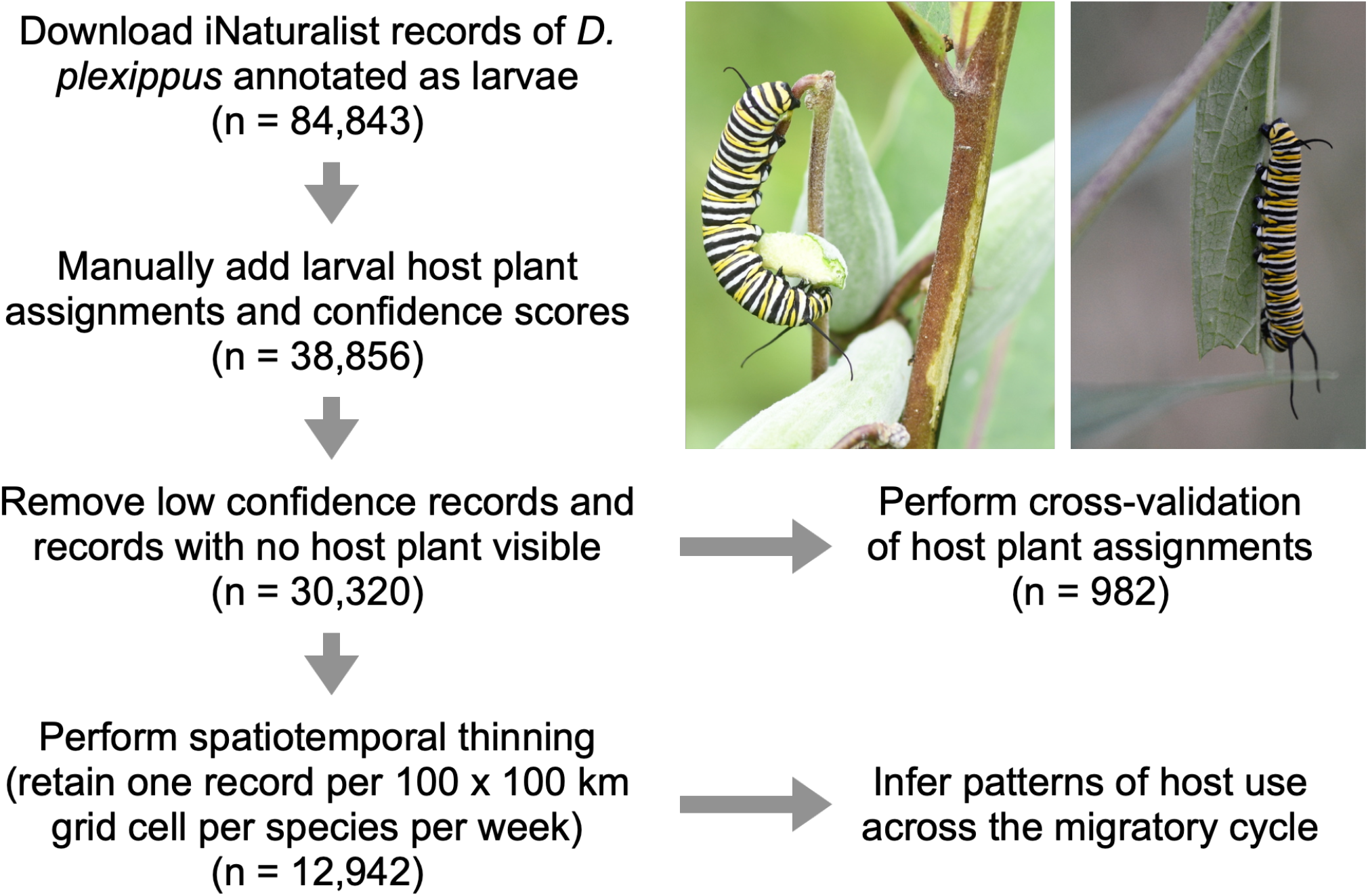
Schematic illustrating data generation and filtering steps used in this project. Left image is an <u>observation</u> of monarch larva on *A. syriaca* in York, ON, Canada; right image is an <u>observation</u> of a monarch larva on *A. incarnata* in Kankakee Co., IL, USA. Image credit: Micah Freedman.

## Results

I manually annotated 38,856 records of monarch larvae from North America (Canada, the United States, and Mexico) with information on host plant identity. After removing low-confidence assignments and records with no host visible, I retained 30,320 records (Figure 1). Among these records, 75 unique host plants were documented, including observations from several previously undescribed hosts (e.g., *Asclepias cryptoceras, A. welshii, A. similis*, and *Oxypetalum coeroleum*) (Figure 2, Table S2), considerably expanding the list of reported hosts for *D. plexippus* (Malcolm & Brower 1989; Borders and Lee-Mäder 2018; also see Greenstein et al. 2022). Of these 75 potential host species, a small number of species comprised a disproportionate number of total records: in eastern North America, 82% of records came from *A. syriaca, A. incarnata, A. curassavica*, and *A. tuberosa*, while in western North America, 85% of records came from *A. curassavica, A. fascicularis*, and *A. speciosa*.

**Figure 2.**
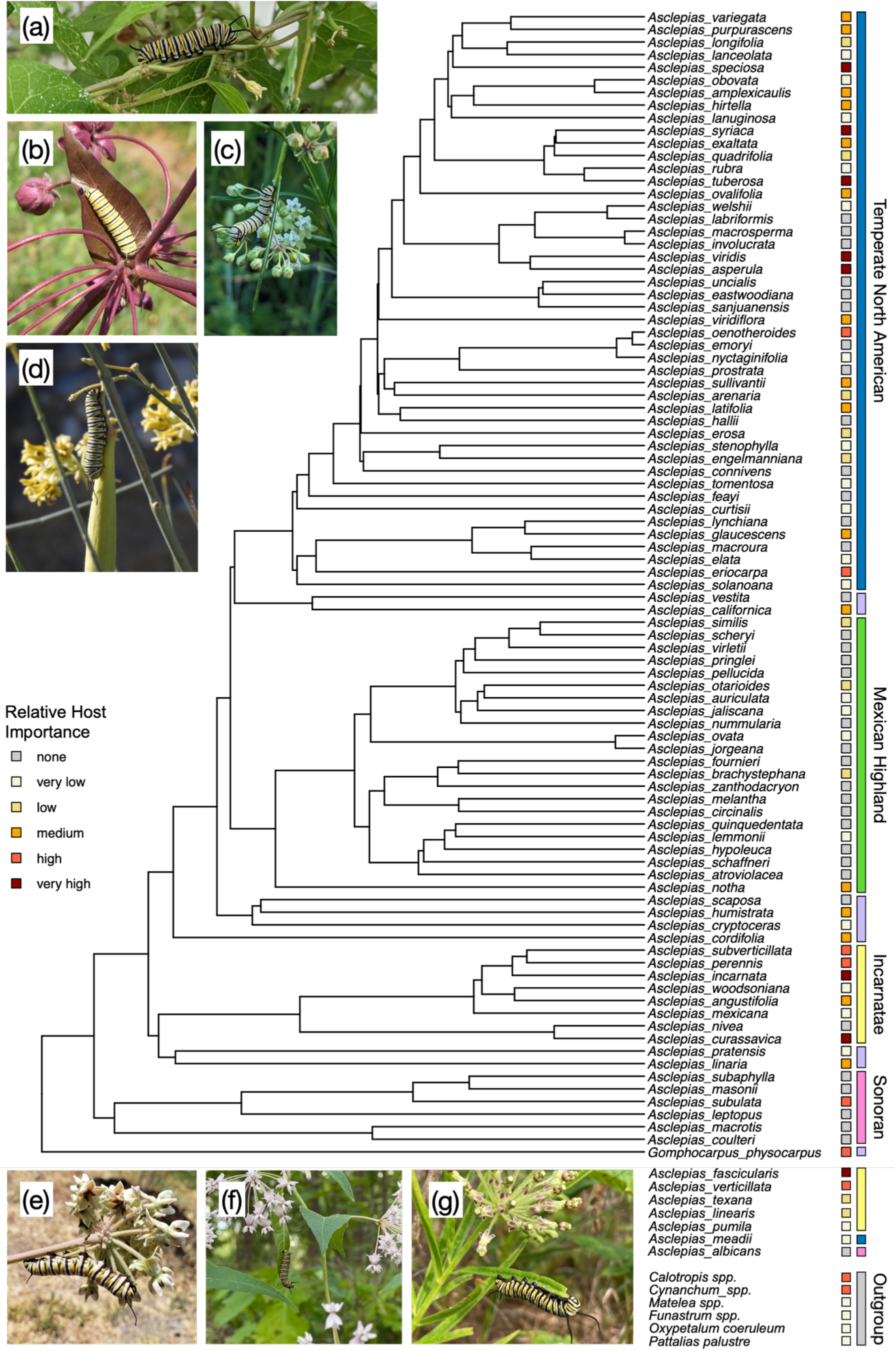
Ultrametric tree of *Asclepias* species and associated clades, based on the phylogeny in Fishbein et al. (2018a). Relative host importance is plotted for each species and corresponds to the number of observed larval records on a particular host (see Methods). Species at the base of the tree were not included in Fishbein et al. (2018a), but their clade-level relationships are known from other work in the genus (e.g., Agrawal and Fishbein 2009). Images highlight the diversity of host species and show (a) *Cynanchum laeve*, <u>Indiana (danlego);</u> (b) *A. cordifolia*, California (<u>paulhavemann</u>); (c) *A. verticillata*, Illinois (<u>allytoth</u>); (d) *A. subulata*, California (dlm1965); (e) *A. glaucescens*, Jalisco (<u>biolcait</u>); (f) *A. quadrifolia*, Missouri (<u>nathanaaron</u>); (g) *A. hirtella*, Illinois (<u>danielpohl</u>).

Based on observed interaction frequency, I categorized milkweed species according to their relative importance as hosts for monarch larvae (Figure 2). As expected, several species from the temperate North American clade were identified as being important hosts, including *A. syriaca, A. tuberosa, A. speciosa, A. viridis, A. asperula, A. oenotheroides*, and *A. eriocarpa*. The Incarnatae clade, despite having relatively few species, featured a high proportion of important hosts, including *A. incarnata, A. fascicularis, A. curassavica, A. perennis, A. verticillata*, and *A. subverticillata*; however, there was no evidence for phylogenetic signal in patterns of host use across the clade (Pagel’s λ = 7.48 × 10^-5^, p = 1.00). I also mapped host use onto the broader phylogeny of the Apocynaceae (Figure 3), confirming that monarch host plants are limited to the “core milkweeds” (Apocynaceae: Asclepiadoideae, formerly treated as Asclepiadaceae), despite evidence for oviposition on host genera outside of this group (Figure 3, Table S3). Interestingly, species in the genus *Vincetoxicum* are not known to support monarch larval development and have been suggested to be “evolutionary traps” for ovipositing monarchs (DiTomasso & Losey 2003, Casagrande & Dacey 2014), despite being nested within a clade of otherwise suitable hosts.

**Figure 3.**
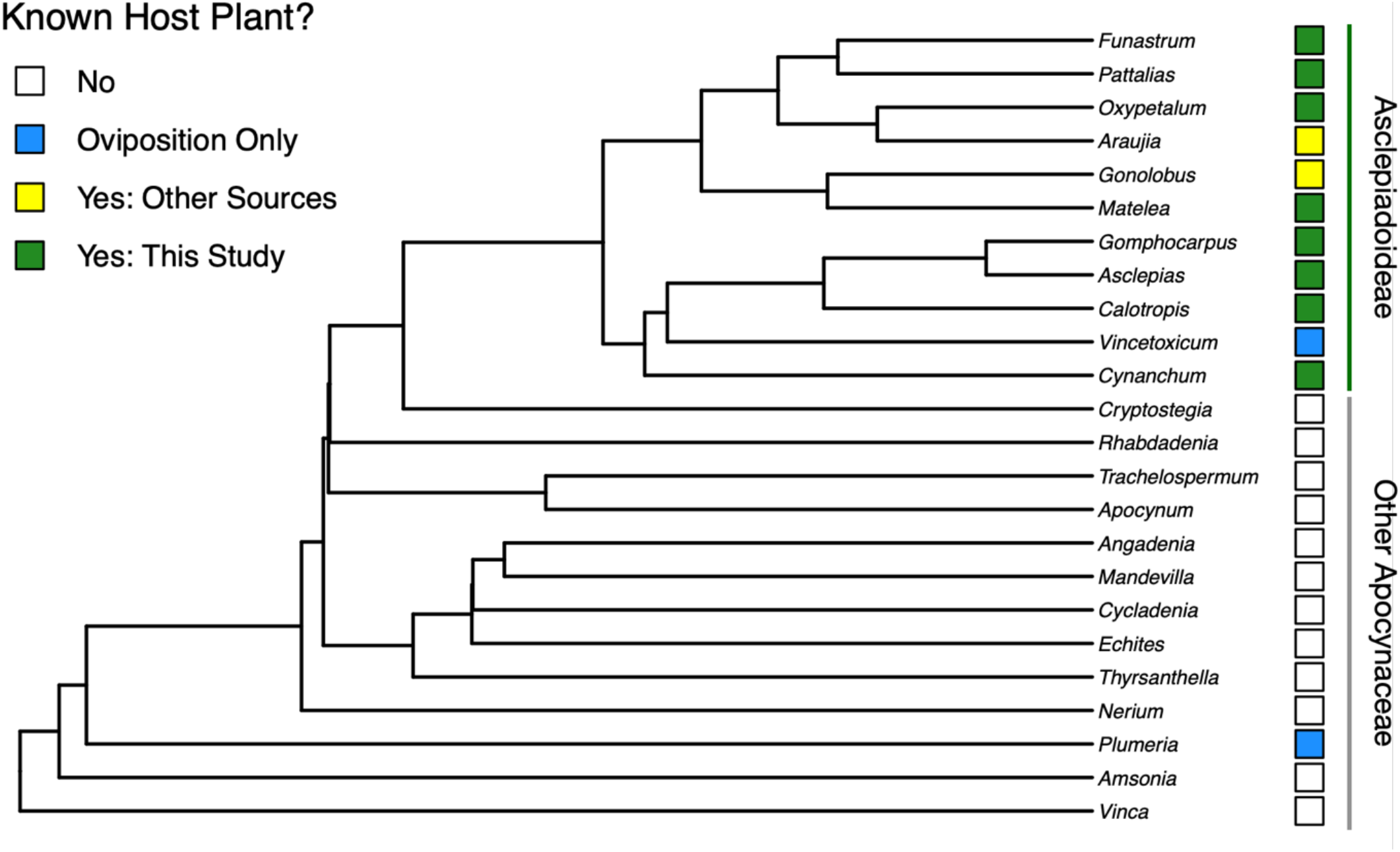
Ultrametric tree depicting selected Apocynaceae genera known to co-occur in North America with monarch butterflies. Host plant status is plotted for each genus, confirming that suitable hosts (i.e., those that are known to support development through adult eclosion) are limited to the core Asclepiadoideae. For details on host plant status, see Methods and Table S3.

I next performed spatial and temporal thinning of records to understand patterns of host use across the monarch’s migratory cycle within North America, retaining a total of 12,942 records. Many North American milkweed species show clear evidence for unimodal peaks of host use during late summer (Figure 4a), which corroborates previously published research suggesting that monarch larvae are generally most abundant in their core summer breeding range in July and August (Prysby & Oberhauser 2004). A handful of milkweed species show evidence for early peaks in records of host use between late April – late May: this applies to *A. viridis, A. asperula, A. amplexicaulis*, and *A. humistrata* in eastern North America and *A. californica* and *A. cordifolia* in western North America (Figure 4b; Malcolm et al. 1987, MacArthur-Waltz et al. 2025). Monarch populations are lowest and may be most demographically sensitive in the first generation that reproduces post-overwintering (Oberhauser et al. 2017), highlighting that these early-season hosts and their degree of phenological overlap with monarchs may play an outsized role in determining monarch abundance in subsequent generations (Ries et al. 2015); the importance of these early-season hosts is also illustrated in Figure 4b. Interestingly, species such as *A. viridis* and *A. oenotheroides* show evidence for bimodal peaks as larval hosts, indicating that monarchs which undergo a round of reproduction in the southern parts of their range in September and October (Flockhart et al. 2013, Fyson et al. 2025) are associated with these hosts.

**Figure 4.**
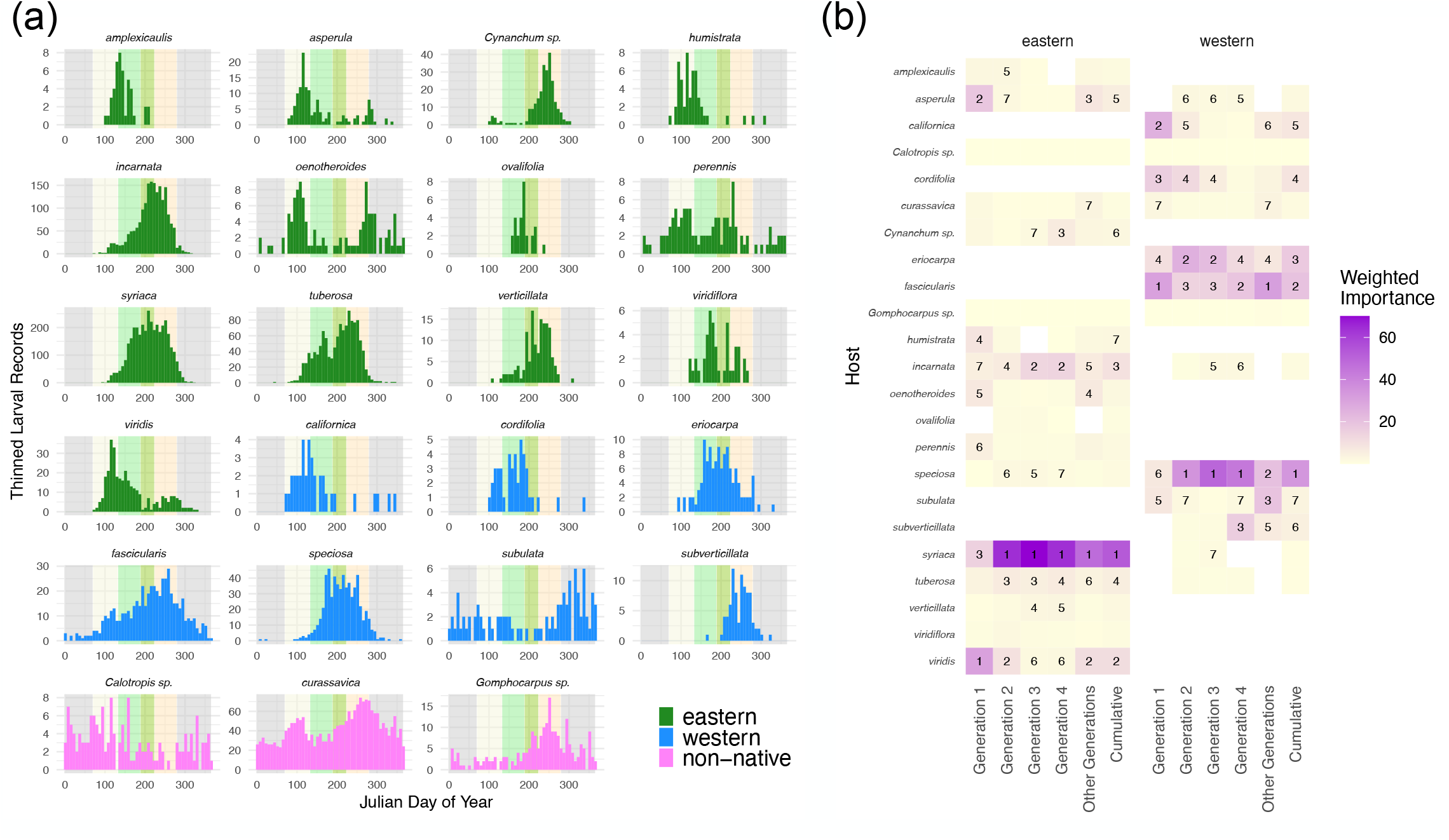
(a) Distribution of host plant use across the year for the 23 most frequently recorded hosts in the dataset. Hosts are separated based on whether their distributions are primarily in eastern or western North America, with hosts not native to temperate North America displayed separately. Counts depicted are based on the thinned dataset (12,942 records) and are grouped into 52 bins, each corresponding to a week of the year. Background coloration within each panel corresponds to the approximate timing of generations within North America (light yellow = Gen1, light green = Gen2, darker green = Gen3, beige = Gen4, grey = other generations). (b) Weighted importance of the same 23 host plants across generations for eastern and western North America. Here, weighted importance is proportional to the probability of a given host species being a wild, non-cultivated plant; hence, species like *A. curassavica* that are predominantly cultivated are down-weighted and receive lower importance scores. Numbers show the rank of the top seven hosts within each generation. For details on weighting, see Methods.

Notably, three milkweed taxa that are not native to temperate areas of North America (*Calotropis spp*., *A. curassavica*, and *Gomphocarpus spp*.), but which have been widely planted horticulturally, supported records of monarch larvae across the entire calendar year (Figure 4a, Figure S1). This pattern is especially pronounced for *A. curassavica*, which represents 64.8% (605/934) of all thinned larval host records in North America between November-February. The spatial extent of winter breeding, as evidenced by records of larvae between November-February, is considerably larger due to non-native hosts: 60% (209/348) of the grid cells that showed evidence for winter breeding featured records exclusively from non-native hosts (Figure 5). While the majority of overall winter-breeding records come from non-native hosts, several native hosts also support winter breeding, including both eastern (*A. oenotheroides, A. perennis*) and western species (*A. fascicularis, A. subulata*). Winter records of larvae on native hosts were very common in western North America: of the 70 western grid cells with records of larvae during winter, 81% (57/70) included records of larvae on native western host plants.

**Figure 5.**
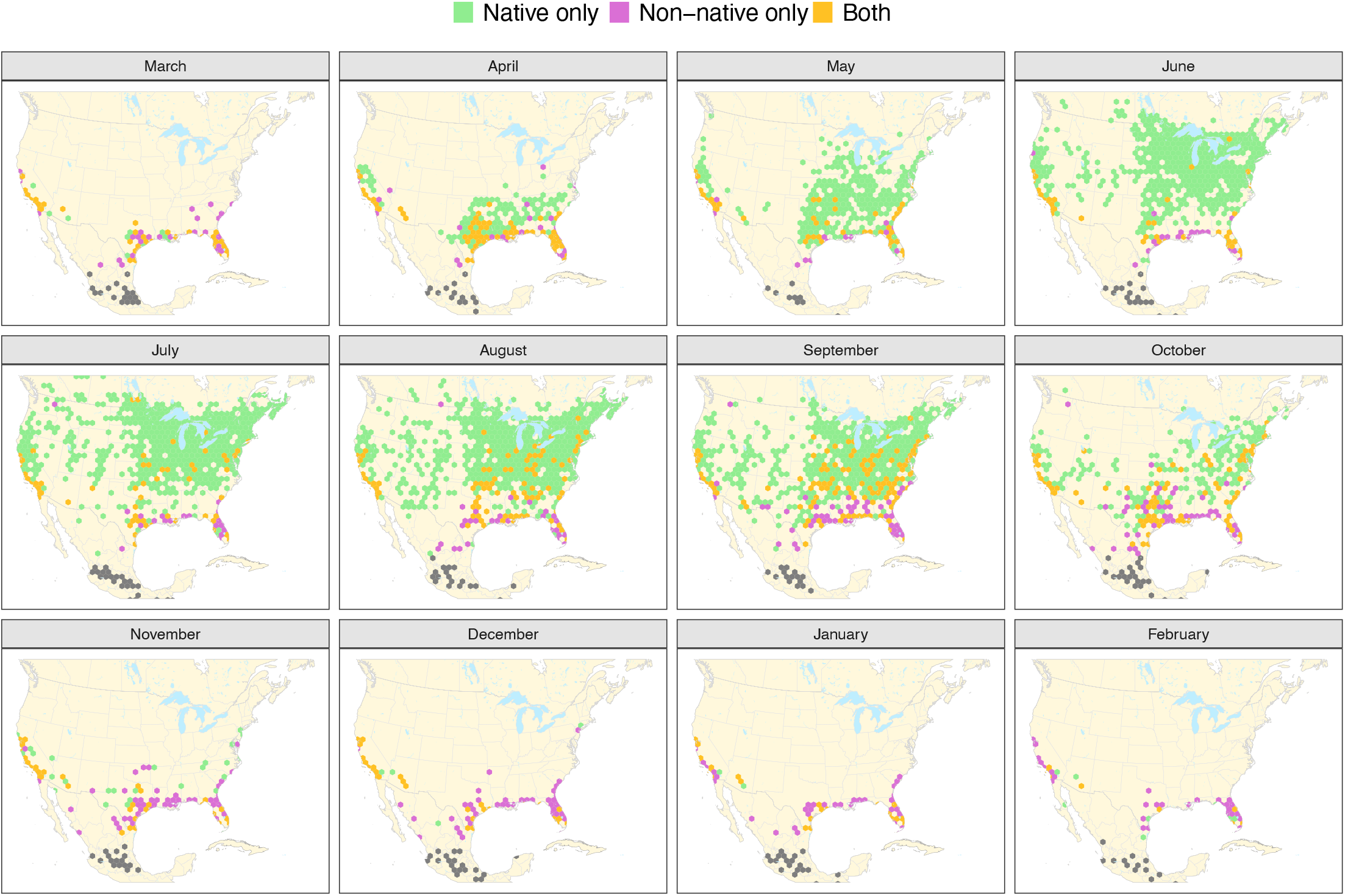
Maps showing monthly distributions of larval records on native vs. non-native host plants. Grid cells are 100 km in diameter and are colored to reflect whether, for a particular month, that cell contained records of larvae on exclusively native hosts (green), exclusively non-native hosts (pink), or a mix of both (orange). Points in grey are records from south of 24ºN latitude, where the native status of *A. curassavica* is ambiguous. Winter records of larvae, in the bottom four panels, show a high fraction of non-native only records, consistent with non-native host plants extending the breeding window for monarchs.

## Discussion

I used more than 35,000 records of monarch butterfly larvae and their associated host plants to document pattern of monarch-milkweed interactions. While using community science images to draw inferences about species interactions is not novel, the number of records reported in this study is approximately 35-fold greater than the mean of studies shown in Table S1, and more than fourfold greater than the next largest study (Santiago-Rosario et al. 2024). This scope of sampling allows for detection of rare host plants (Figure 2), which can be important for understanding the combinations of traits that define host breadth in insect herbivores (Fordyce et al. 2016), as well comprehensive characterization of seasonal patterns of host use across the migratory cycle (Figures 4 and 5).

A small number of species comprised a high fraction of the total number of observed interactions. Perhaps not surprisingly, these are hosts that are geographically widespread and available during periods of the year when monarch larvae are most abundant; indeed, there is a strong positive correlation between extent of occurrence and number of larval records across milkweed species (Figure S2). The high frequency of interactions with this small subset of species should exert strong natural selection on larval performance on these hosts, as has been suggested for *A. syriaca* (Agrawal et al. 2015). The most frequently used hosts are spread across the *Asclepias* phylogeny (Figure 2), suggesting that patterns of spatial and temporal overlap between monarchs and their milkweed hosts, rather than trait similarity among hosts (e.g., Pearse & Hipp 2009), is the major driver of host use frequency in the monarch-milkweed system.

Despite having relatively few overall records, a handful of spring-emergent milkweed species have high weighted importance values as hosts in both eastern and western North America (Figure 4b). Eastern species *A. viridis* and *A. asperula* and western species *A. californica* and *A. cordifolia* were among the top five most important hosts in their respective regions. The prevalence of *A. viridis* as an early-season host is supported by chemical fingerprinting approaches (Malcolm et al. 1989), although no comparable data currently exist for western North America. Surprisingly, *A. syriaca* was identified as the third most important eastern host even in Generation 1 (March 12 – May 12), highlighting that this widespread host emerges and supports monarch larvae quite early in the year in the southern parts of its range (Figure S3).

Three species of milkweed not native to temperate North America (*A. curassavica, C. procera*, and *G. physocarpus*) represented a considerable fraction of overall records, particularly during winter months (Figure 5). Numerous studies have suggested that these hosts might alter monarch life history, either by causing migratory monarchs that encounter them to break reproductive diapause (Batalden & Oberhauser 2015, Majewska & Altizer 2019; but see Fedorka & Rich 2026), or because their atypical phenology and lack of senescence prolongs the temporal window over which larval development is possible (Satterfield et al. 2015, Satterfield et al. 2018a). In the case of *A. curassavica*, clear seasonal peaks of host use that coincide with spring and fall migration are apparent (Figure 4), highlighting the role that this species may play in facilitating interactions between migratory and year-round breeding monarchs (Satterfield et al. 2018b). While this study is certainly not unique in suggesting that non-native milkweeds may alter the breeding niche of monarch butterflies, it does quantify the spatial extent of this phenomenon and illustrates that a substantial fraction of community science records of monarch-milkweed interactions feature cultivated, non-native hosts. Among the more than 16,000 unique observers who contributed to this dataset, around 20% posted records exclusively from non-native hosts, and around 33% posted records exclusively from plants that were likely cultivated (Table S4); this finding highlights that many community scientists’ perceptions of monarch-milkweed interactions may be shaped by observations in “contrived” settings.

Interestingly, several native milkweed species also supported larvae during November-February. Along the U.S. Gulf Coast, *A. oenotheroides* and *A. perennis* hosted larvae throughout the winter, although considerably less frequently than non-native hosts. Winter records of larvae on *A. perennis* are consistent with chemical fingerprinting data that find moderate to high proportions of this host species among winter-active monarchs in northern and central Florida (Moranz and Freedman, unpublished data). In California, *A. fascicularis* also supported year-round breeding (Figure 3) at a spatial scale comparable to *A. curassavica* (Figure S4). More than 58% of larval records on A. *fascicularis* appear to be cultivated plants (Table S4), highlighting the possibility that cultivation status (i.e., plants that receive water/nutrient amendments and remain vegetative year-round versus those that are wild and naturally senesce) may be a more important distinction than native versus non-native status in predicting winter breeding among monarchs (Lee and Freedman, in prep). Human cultural practices that promote atypical milkweed phenology may thus be analogous to other food subsidies that alter animal movement patterns (Satterfield et al. 2018a), such as agricultural waste grain for whooping cranes (Teitelbaum et al. 2016) and garbage dumps for white storks (Gilbert et al. 2016).

A major shortcoming of data sources like iNaturalist is observer bias (Di Cecco et al. 2021): human observers are much more likely to post observations from populated areas, affluent areas, and along roads and hiking trails (Geurts et al. 2023, Della Rocca et al. 2024). Even though spatial and temporal thinning of records can help to mitigate the influence of factors like increased observer activity in urban areas, the inferences drawn in this study are still very unlikely to be fully representative. For example, the species of milkweed from the Mexican highland clade of species depicted in Figure 2 as having little to no importance as hosts may in fact be widely used, but simply are not observed by users of the iNaturalist platform. That said, many of the findings reported here can be corroborated with alternate methods. For example, the observation that *A. syriaca* is the most prevalent host plant used by monarchs in North America is supported by chemical fingerprinting data (Seiber et al. 1986, Malcolm et al. 1989) and population genomic evidence, which points to its shared demographic history with monarchs (Boyle et al. 2023). Ongoing chemical fingerprinting work, on-the-ground surveys, and the ever-increasing spatial extent of iNaturalist observations will continue to illustrate the extent to which these data are representative of true patterns of host use.

Community science platforms will continue to play an important role in biodiversity research in the 21^st^ century: iNaturalist recently reached 300 million total observations and is on pace to surpass the estimated 1.1 billion specimens in natural history museums around the world by 2040. Datasets such as the one presented here highlight the value of community science records and can also be used as training data for automated computer vision approaches that make documentation of species interactions higher throughput (Lürig et al. 2021). Species interactions can be challenging to document at scale, and the prospect of their “insidious extinction” (Janzen 1974) emphasizes the importance of utilizing all available data sources to record them in the face of global change.

## Methods

### Data Sourcing, Record Annotation, and Validation

I used the iNaturalist API to download all “Research Grade” records of monarch larvae from Canada, the United States, and Mexico; records were downloaded several times between 2022-2025, with the most recent update on 29-July-2025. This resulted in a total of 84,843 larval records. Improper identifications of monarch larvae were rare (<1% of records, pers. obs.), with most misidentifications featuring queens (*D. gillipus*), milkweed tussock moths (*Euchaetes egle*), or hooded owlets (*Cucullia spp*.). Host plant annotations were added to a subset of records (n = 38,856) in a semi-randomized fashion. First, I used a random number generator to create four-digit values and then annotated all records whose URLs contained that value. Second, for states and provinces with sparse records or locations that were deemed more likely to contain “interesting” host plant records, I added annotations to all possible records. Finally, to maximize the spatial coverage of records in the database, I generated pairwise distance matrices of all records within each state/province and annotated any record located >10 km from the nearest other larval record. I included records with obscured coordinates, as the degree of location uncertainty for these records is relatively small compared to the scale at which I conducted spatial thinning (see below).

Host plant annotations were added to each larval record manually and were made based on (1) visible features of the plant shown in the observation, which were checked against the characters outlined in the monograph of *Asclepias* (Woodson 1954); (2) corresponding Research Grade observations of milkweed plants by the same observer on the same day and in the same location; (3) prior knowledge of the geographic ranges of each North American milkweed species; (4) comments from the observer associated with the observation. For each of the 38,856 annotated observations, I gave a score of “certain,” “high,” “medium,” or “low” to reflect confidence in the host plant assignment; records with no host plant visible (n = 3,641) were assigned as “unknown.” For all analyses, only records with confidence scores of “certain,” “high,” or “medium” were used (n = 30,320); this comprised the core unfiltered dataset. In areas where *A. syriaca* and *A. speciosa* are sympatric, leaf characters alone are not sufficient to distinguish the species; as such, these sympatric records (n = 623) were recorded as “complex speciosa/syriaca” and not included in subsequent analyses.

To determine whether host plant assignments were reliable, I selected approximately 3% of records from the core unfiltered dataset (n = 982) for cross-validation using the computer-vision-based plant identification app PictureThis®. Records from each of the 36 most commonly observed hosts were chosen for cross-validation, with greater numbers of records chosen for species that were observed more commonly (Table S5). After removing taxa that were apparently not included in the PictureThis® species database (i.e., cross-validation accuracy of 0%, even when tested against high-quality default taxon photos), records from the remaining 30 species had overall cross-validation accuracies of 94.7%; notably, this included accuracies of 100% for *A. tuberosa* (62/62), 100% for A. *incarnata* (68/68), 96.6% for *A. syriaca* (114/118), 94.6% for *A. viridis* (35/37), and 92.6% for *A. curassavica* (88/95). Thus, overall, I feel confident in the host plant assignments that comprise this dataset.

For some host taxa (e.g., *Cynanchum*), I recorded only the genus and did not attempt species-level identifications for all observations. Thus, records for some taxa such as *Calotropis spp*. reflect a mix of *C. procera* and *C. gigantea*. For the full list of 75 species recorded as hosts in this dataset, see Table S2. In addition to assignments of host plant identity, I also included annotations of whether the host plant was cultivated.

### Host Use Across the Milkweed Phylogeny

To visualize patterns of host use across the *Asclepias* phylogeny, I used the whole-plastome tree file from Fishbein et al. (2018a) and plotted it as an ultrametric tree. For taxa not included in the whole-plastome tree but present in the host use dataset, I used previously published phylogenies of the genus to assign these remaining taxa to respective clades. I scored relative host importance as a function of the number of thinned records in the dataset: none = 0 records, very low = 1-5 records, low = 6-20 records, medium = 21-100 records, high = 101-500 records, very high = >500 records. To test for phylogenetic signal in patterns of host use, I calculated Pagel’s λ using the phylosig() function from phytools v. 2.0 (Revell 2024).

To visualize broader patterns of host use across the Apocynaceae, I used the nuclear tree from Fishbein et al. (2018b) and pruned it to include representative species from 11 genera within the core Asclepiadoideae and 14 additional genera from the family. All genera displayed in Figure 3 are present in North America and are potential hosts, although several are represented only as non-native ornamental taxa. Some taxa have been reported as suitable hosts for larval development but were not recorded as hosts in this dataset; similarly, some taxa have records of monarch oviposition but are not known to support larval development (see Table S3 for details).

### Thinning of Records and Weighted Host Importance

Because iNaturalist observations are generated in a highly non-random way, I performed spatial and temporal thinning prior to conducting analyses related to patterns of host use. I defined a mesh of hexagonal grid cells (hexgrids) with a diameter of 100 km. I then used the ordinal date of observation to determine the week of the year for each record, generating an integer value of 1-52 for each record. For each combination of hexgrid ID *x* week *x* host species, I retained a single record for use in downstream analysis, resulting in 12,982 thinned records. I treated *A. curassavica, Calotropis spp*., and *Gomphocarpus spp*. as non-native hosts; while this assumption is appropriate for the latter two taxa, the “ancestral” range of *A. curassavica* is less clear, though has been suggested to be Central America, South America, and the Caribbean (Woodson 1954). Thus, in Figure 5 and associated analyses, I considered all records from 24º of latitude and further south to be ambiguous in terms of their native status.

To determine weighted importance of host species (Figure 4B), I considered only the 23 most abundant species in the thinned dataset. For each host species, I used the sum of records whose cultivation status was listed as “no” or “unlikely” to calculate the proportion of non-cultivated records. Next, I partitioned records based on their approximate generation (Generation 1: March 12 – May 12; Generation 2: May 13 – July 7; Generation 3: July 8 – Aug 11; Generation 4: Aug 12 – October 6; Other Generations: all remaining dates). Within each generation, I calculated weighted importance as the proportion of total records associated with a particular host multiplied by the proportion of that host’s records from non-cultivated plants. Here, the rationale is to further account for observer bias by assuming that observers are much more likely to post records from easily accessible locations (e.g., backyards, parks, cities) that contain cultivated plants, and assign greater weight to records from areas with naturally growing hosts. Finally, to calculate cumulative weighted importance, I summed the weighted importance across generations; for records in the category “Other Generations,” I multiplied their relative importance by 0.1 to reflect that a relatively small proportion of monarchs likely engage in breeding outside of the primary four generations. For alternative formulations of host importance that use different assumptions, see Figure S5.

## References

1. A. A. Agrawal, et al., Phylogenetic ecology of leaf surface traits in the milkweeds (Asclepias spp.): chemistry, ecophysiology, and insect behavior. New Phytol. 183, 848–867 (2009).

2. A. A. Agrawal, J. G. Ali, S. Rasmann, M. Fishbein, Macroevolutionary trends in the defense of milkweeds against monarchs. In: Monarchs in a changing world: Biology and conservation of an iconic butterfly 47–59 (2015).

3. R. V. Batalden, K. S. Oberhauser, Potential changes in eastern North American monarch migration in response to an introduced milkweed, Asclepias curassavica. In: Monarchs in a changing world: Biology and conservation of an iconic butterfly 215–224 (2015).

4. J. H. Boyle, et al., Temporal matches between monarch butterfly and milkweed population changes over the past 25,000 years. Curr. Biol. 33, 3702-3710.e5 (2023).

5. C. T. Callaghan, et al., Three frontiers for the future of biodiversity research using citizen science data. BioScience 71, 55–63 (2021).

6. M. D. de Groot, et al., Beetlehangers.org: harmonizing host–parasite records of Harmonia axyridis and Hesperomyces harmoniae. Arthropod Plant Interact. 18, 665–679 (2024).

7. F. Della Rocca, M. Musiani, M. Galaverni, P. Milanesi, Improving online citizen science platforms for biodiversity monitoring. J. Biogeogr. 51, 2412–2423 (2024).

8. G. J. Di Cecco, et al., Observing the observers: How participants contribute data to iNaturalist and implications for biodiversity science. Bioscience 71, 1179–1188 (2021).

9. A. DiTommaso, J. E. Losey, Oviposition preference and larval performance of monarch butterflies (Danaus plexippus) on two invasive swallow-wort species: Monarch butterflies prefer milkweed over two swallow-worts. Entomol. Exp. Appl. 108, 205–209 (2003).

10. B. Borders, E. Lee-Mäder, Milkweeds: A conservation practitioners guide. The Xerces Society for Invertebrate Conservation. (2018).

11. K. M. Fedorka, M. Rich, The effect of temperature and tropical milkweed on monarch butterfly migration disruption. R. Soc. Open Sci. 13, 251537 (2026).

12. M. Fishbein, et al., Evolution at the tips: Asclepias phylogenomics and new perspectives on leaf surfaces. Am. J. Bot. 105, 514–524 (2018).

13. M. Fishbein, et al., Evolution on the backbone: Apocynaceae phylogenomics and new perspectives on growth forms, flowers, and fruits. Am. J. Bot. 105, 495–513 (2018).

14. D. T. T. Flockhart, et al., Tracking multi-generational colonization of the breeding grounds by monarch butterflies in eastern North America. Proc. Biol. Sci. 280, 20131087 (2013).

15. J. A. Fordyce, C. C. Nice, C. A. Hamm, M. L. Forister, Quantifying diet breadth through ordination of host association. Ecology 97, 842–849 (2016).

16. C. J. L. Fourreau, et al., The Trojan seahorse: citizen science pictures of a seahorse harbour insights into the distribution and behaviour of a long-overlooked polychaete worm. Proc. Biol. Sci. 291, 20241780 (2024).

17. V. K. Fyson, G. W. Mitchell, J. F. Wilmshurst, C. Callaghan, Mapping monarch seasonal breeding patterns in Eastern North America to inform mowing strategies for roadsides and other rights-of-ways. J. Insect Conserv. 29, 41 (2025).

18. E. M. Geurts, J. D. Reynolds, B. M. Starzomski, Not all who wander are lost: Trail bias in community science. PLoS One 18, e0287150 (2023).

19. N. I. Gilbert, et al., Are white storks addicted to junk food? Impacts of landfill use on the movement and behaviour of resident white storks (Ciconia ciconia) from a partially migratory population. Mov Ecol 4, 7 (2016).

20. L. Greenstein, C. Steele, C. M. Taylor, Host plant specificity of the monarch butterfly Danaus plexippus: A systematic review and meta-analysis. PLoS One 17, e0269701 (2022).

21. N. Jansen, N. Pruijn, M. Mayer, Citizen observations shed new light on geographic variation in colour polymorphism of a widespread reptile. J. Biogeogr. 52, 629–640 (2025).

22. A. Johnston, et al., Analytical guidelines to increase the value of community science data: An example using eBird data to estimate species distributions. Divers. Distrib. 27, 1265–1277 (2021).

23. M. D. Lürig, S. Donoughe, E. I. Svensson, A. Porto, M. Tsuboi, Computer vision, machine learning, and the promise of phenomics in ecology and evolutionary biology. Front. Ecol. Evol. 9 (2021).

24. D. J. MacArthur-Waltz, D. S. Sommer, P. Singh, L. H. Yang, Temperature and precipitation explain species-specific phenological patterns in five native California milkweed species (Asclepias spp.). J. Ecol. 113, 3637–3649 (2025).

25. A. A. Majewska, S. Altizer, Exposure to Non-Native Tropical Milkweed Promotes Reproductive Development in Migratory Monarch Butterflies. Insects 10 (2019).

26. S. B. Malcolm, L. P. Brower, Evolutionary and ecological implications of cardenolide sequestration in the monarch butterfly. Experientia (1989).

27. S. B. Malcolm, B. J. Cockrell, L. P. Brower, Monarch Butterfly Voltinism: Effects of Temperature Constraints at Different Latitudes. Oikos 49, 77–82 (1987).

28. S. B. Malcolm, B. J. Cockrell, L. P. Brower, Cardenolide fingerprint of monarch butterflies reared on common milkweed, Asclepias syriaca L. J. Chem. Ecol. 15, 819–853 (1989).

29. B. M. Mason, et al., iNaturalist accelerates biodiversity research. Bioscience 75, 953–965 (2025).

30. P. F. McKenzie, A. E. Berardi, R. Hopkins, Delayed flowering phenology of red-flowering plants in response to hummingbird migration. Curr. Biol. 35, 2175-2182.e3 (2025).

31. K. Oberhauser, et al., A trans-national monarch butterfly population model and implications for regional conservation priorities: Regional monarch conservation priorities. Ecol. Entomol. 42, 51–60 (2017).

32. R.J. Ortíz-Zárate, et al., Bot fly parasitism in mantled howler monkeys (Alouatta palliata): General patterns and climate influences. Am. J. Primatol. 86, e23680 (2024).

33. I. S. Pearse, A. L. Hipp, Phylogenetic and trait similarity to a native species predict herbivory on non-native oaks. Proc. Natl. Acad. Sci. U. S. A. 106, 18097–18102 (2009).

34. N. Pernat, et al., Extracting secondary data from citizen science images reveals host flower preferences of the Mexican grass-carrying wasp Isodontia mexicana in its native and introduced ranges. Ecol. Evol. 14, e11537 (2024).

35. M. D. Prysby, K. S. Oberhauser, Temporal and geographic variation in monarch densities: Citizen Scientists Document Monarch Population Patterns. Pages 9-20. Monarch Butterfly Biol Conserv (2004).

36. B. J. Putman, R. Williams, E. Li, G. B. Pauly, The power of community science to quantify ecological interactions in cities. Sci. Rep. 11, 3069 (2021).

37. L. J. Revell, phytools 2.0: an updated R ecosystem for phylogenetic comparative methods (and other things). PeerJ 12, e16505 (2024).

38. L. Ries, D. J. Taron, E. Rendón-Salinas, K. S. Oberhauser, Connecting Eastern Monarch Population Dynamics across Their Migratory Cycle. In: Monarchs in a changing world: Biology and conservation of an iconic butterfly 47–59 (2015).

39. L. Ries, K. Oberhauser, A citizen army for science: Quantifying the contributions of citizen scientists to our understanding of monarch butterfly biology. Bioscience 65, 419–430 (2015).

40. R. M. Rosa, D. C. Cavallari, R. B. Salvador, iNaturalist as a tool in the study of tropical molluscs. PLoS One 17, e0268048 (2022).

41. L. Y. Santiago-Rosario, J. Book, S. Mathews, Continental-scale interactions of Australian showy mistletoes and their hosts. Am. J. Bot. 111, e16443 (2024).

42. D. A. Satterfield, P. P. Marra, T. S. Sillett, S. Altizer, Responses of migratory species and their pathogens to supplemental feeding. Philos. Trans. R. Soc. Lond. B Biol. Sci. 373, 20170094 (2018).

43. D. A. Satterfield, J. C. Maerz, S. Altizer, Loss of migratory behaviour increases infection risk for a butterfly host. Proc. Biol. Sci. 282, 20141734 (2015).

44. D. A. Satterfield, et al., Migratory monarchs that encounter resident monarchs show life-history differences and higher rates of parasite infection. Ecol. Lett. 21, 1670– 1680 (2018).

45. J. N. Seiber, et al., Cardenolide connection between overwintering monarch butterflies from Mexico and their larval food plant, Asclepias syriaca. Journal of Chemical Ecology (1986).

46. P. A. Silva, G. N. da Silva Júnior, L. S. Santos, L. Brito, Revealing new insights into Red-bellied Macaw foraging ecology through citizen photography. Austral Ecol. 49 (2024).

47. C. S. Teitelbaum, et al., Experience drives innovation of new migration patterns of whooping cranes in response to global change. Nat. Commun. 7, 12793 (2016).

48. J. C. Tourville, G. L. D. Murray, S. J. Nelson, Distinct latitudinal patterns of shifting spring phenology across the Appalachian Trail Corridor. Ecology 105, e4403 (2024).

49. R. Vardi, O. Berger-Tal, U. Roll, iNaturalist insights illuminate COVID-19 effects on large mammals in urban centers. Biol. Conserv. 254, 108953 (2021).

50. R. E. Woodson, The North American Species of Asclepias L. Ann. Mo. Bot. Gard. 41, 1–211 (1954).

