## Supplementary figures and images for "Patterns of host plant use by monarch butterflies revealed through annotation of more than 35,000 community science records"

### Figure S1 - pooled histograms.pdf

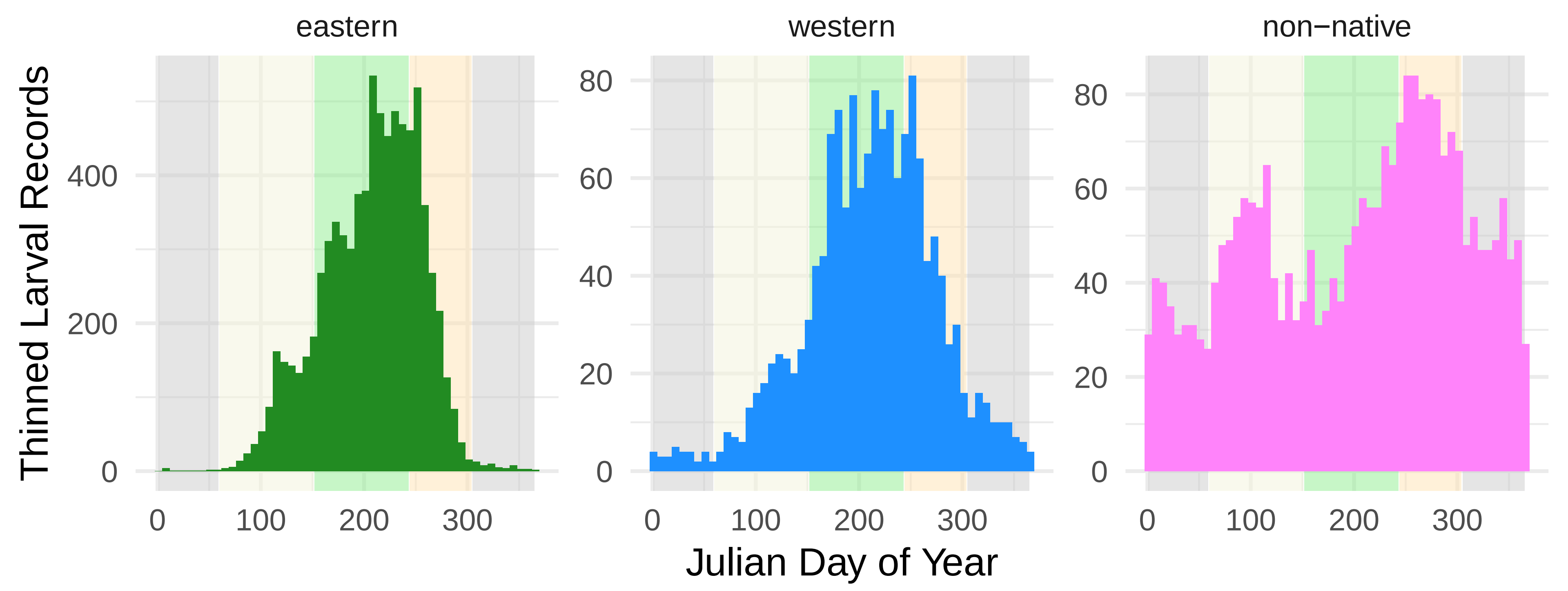

### Figure S2 - range size.pdf

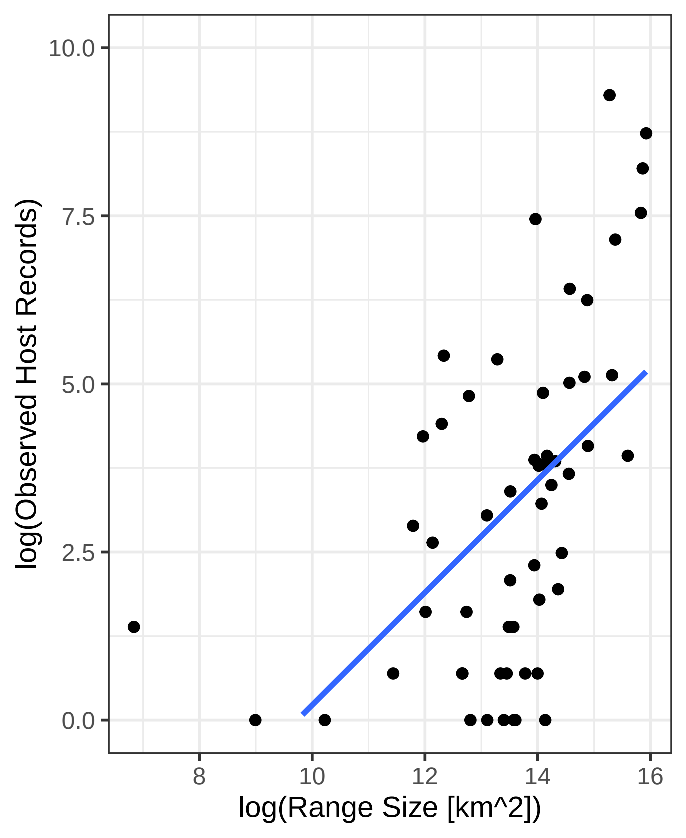

### Figure S3 - syriaca.pdf

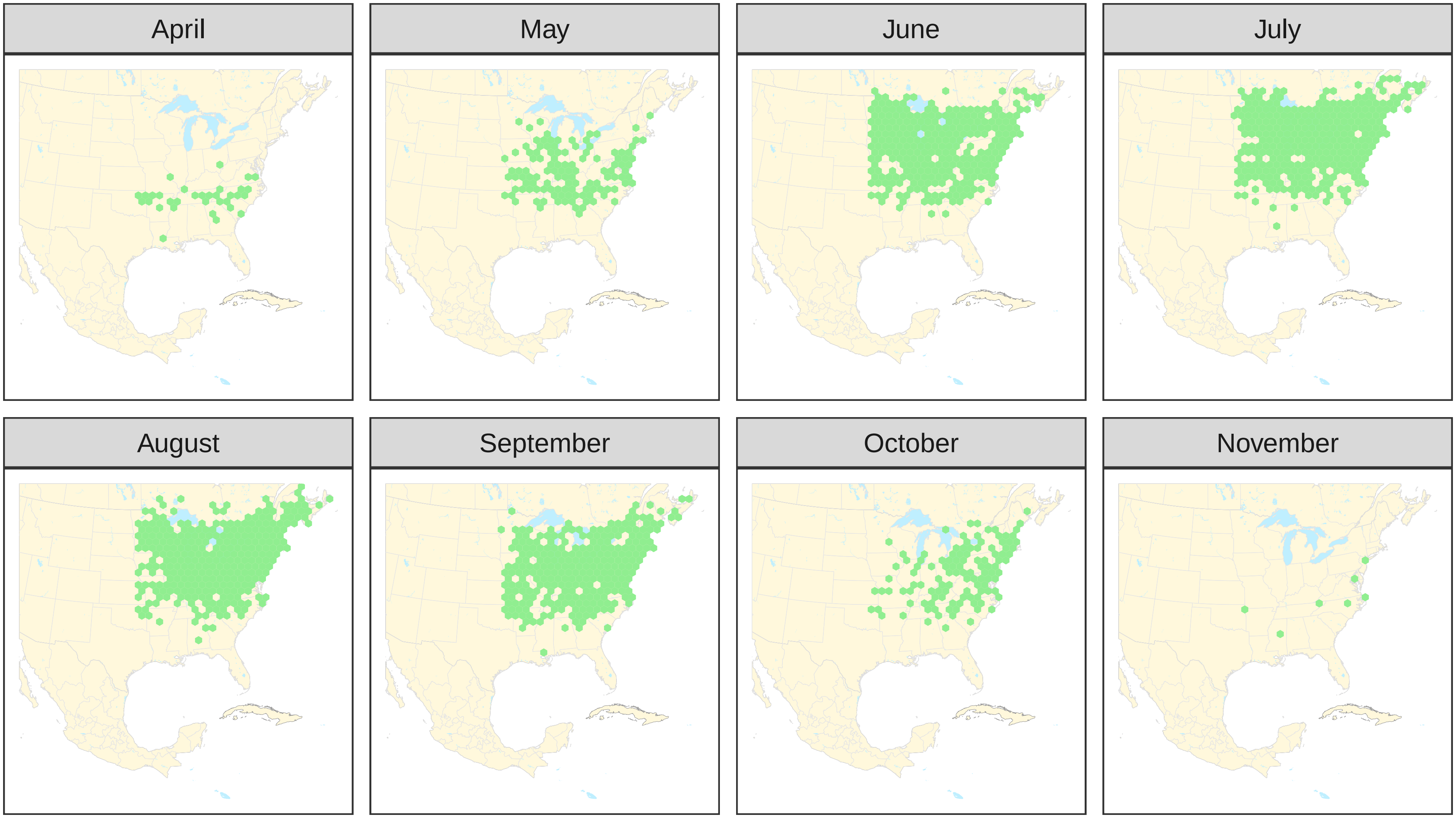

### Figure S4 - fasc and cur.pdf

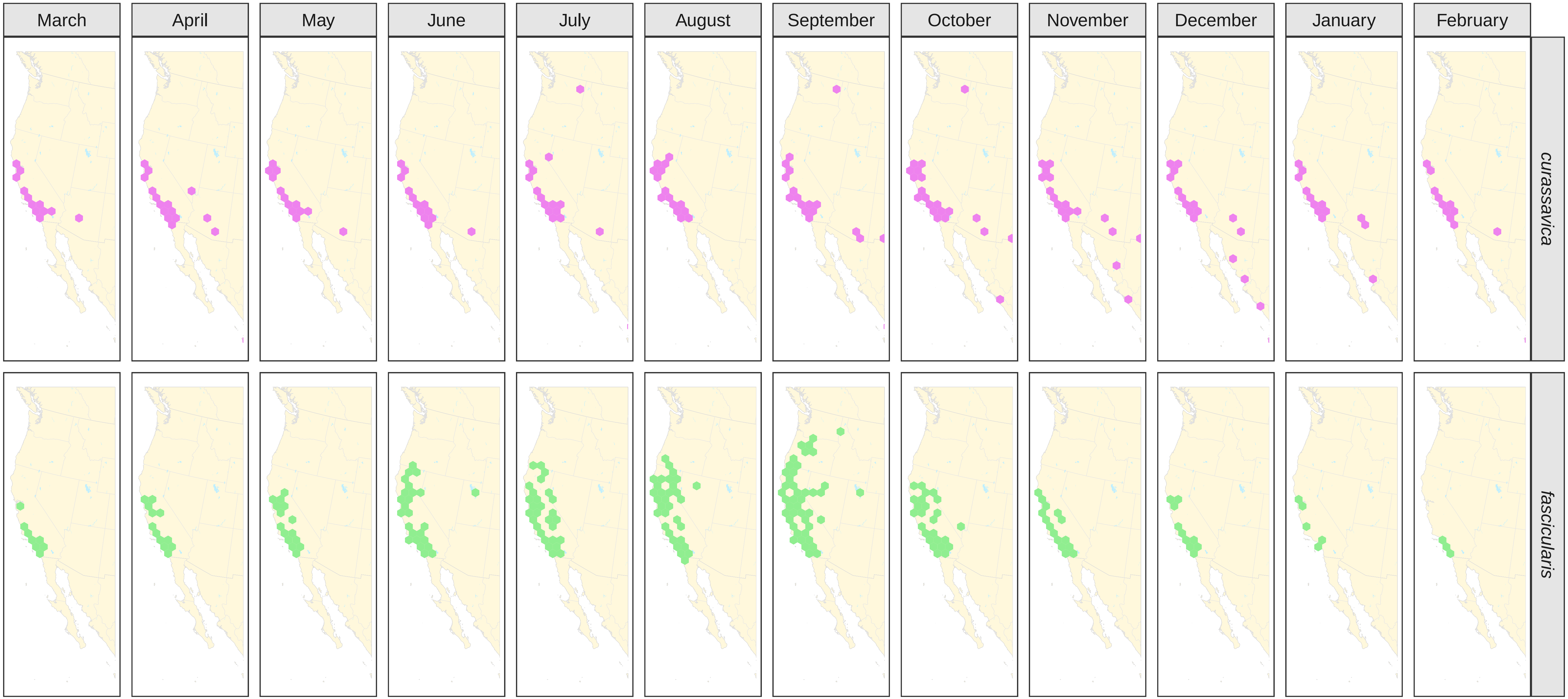

### Figure S5 - alternative weighting.pdf

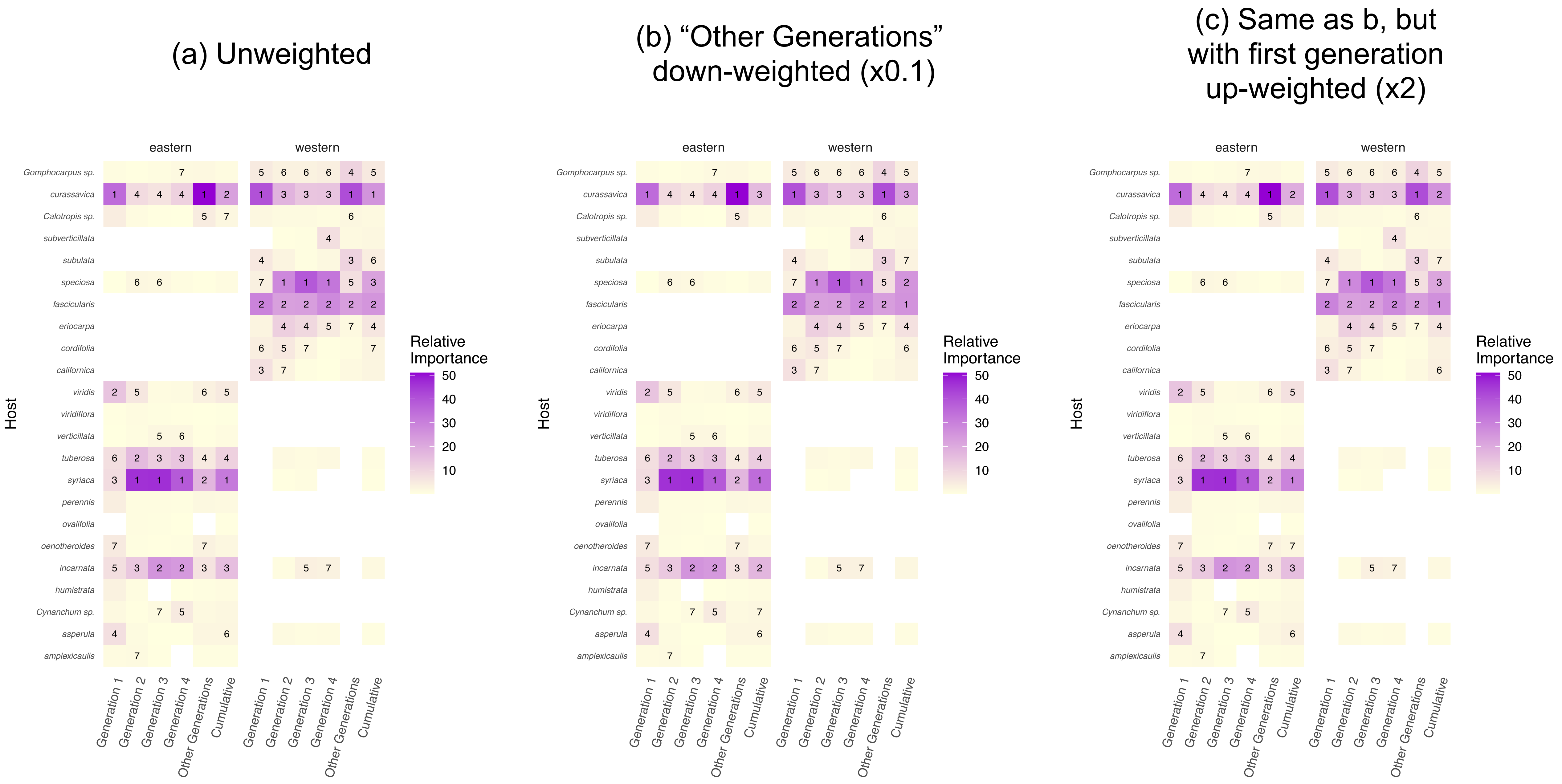
